# Fitness effects of thermal stress differ between outcrossing and selfing populations in *Caenorhabditis elegans*

**DOI:** 10.1101/074328

**Authors:** Agata Plesnar-Bielak, Marta K. Labocha, Paulina Kosztyła, Katarzyna R. Woch, Woronika M. Banot, Karolina Markot, Magdalena Skarboń, Monika A. Prus, Zofia M. Prokop

## Abstract

The maintenance of males and outcrossing is widespread, despite considerable costs of males. By enabling recombination between distinct genotypes, outcrossing may be advantageous during adaptation to a novel environments and if so, it should be selected for under environmental challenge. However, a given environmental change may influence fitness of male, female, and hermaphrodite or asexual individuals differently, and hence the relationship between reproductive system and dynamics of adaptation to novel conditions may not be driven solely by the level of outcrossing and recombination. This has important implications for studies investigating the evolution of reproductive modes in the context of environmental changes, and for the extent to which their findings can be generalized. Here, we use *Caenorhabditis elegans* – a free-living nematode species in which hermaphrodites (capable of selfing but not cross-fertilizing each other) coexist with males (capable of fertilizing hermaphrodites) – to investigate the response of wild type as well as obligatorily outcrossing and obligatorily selfing lines to stressfully increased ambient temperature. We found that thermal stress affects fitness of outcrossers much more drastically than that of selfers. This shows that apart from the potential for recombination, the selective pressures imposed by the same environmental change can differ between populations expressing different reproductive systems and affect their adaptive potential.

## Introduction

Outcrossing, *i.e.* reproduction by fusing gametes of distinct individuals, remains one of evolution’s mysteries. Compared to uniparental reproduction (asexuality or self-fertilization), it incurs considerable costs, particularly when associated with the production of males which facilitate outcrossing but do not themselves bear offspring, while requiring energy resources that could have been used otherwise (Maynard Smith 1971; 1978; Lloyd 1980; Bell 1982; Uyenoyama 1984; Lively and Lloyd 1990; Anderson et al. 2010). Nevertheless, a vast majority of animal species produce males, suggesting that this mode of reproduction does bring some significant selective advantages. Most theoretical explanations proposed to date relate to the role of recombination. Outcrossing shuffles genes among individuals, creating new combinations of alleles. Therefore, it can break apart selection interference between beneficial and deleterious mutations (Hill-Robertson effect), facilitating the spread of the former and purging of the latter (reviewed by Otto 2009). Importantly, this may also lead to (some of) the offspring of outcrossing individuals having increased fitness in a changing environment (Stebbins 1957). Both these factors can accelerate the rate of adaptation to novel environmental conditions. Thus, the benefits of outcrossing should be particularly pronounced under environmental change (e.g. Colegrave 2002; Goddard et al. 2005; Morran et al. 2009, but see Zeyl & Graham Bell 1997).

However, selective pressures imposed by a given environmental factor might differ between reproductive modes. In other words, the same change in the external environment may differently influence the fitness of dioecious and hermaphrodite or asexual individuals of the same species, such that fitness differences in adapting populations might not result solely from the differences in outcrossing rates. Furthermore, these effects may be specific to environmental conditions under study. If the novel environment applied in a study is more stressful for hermaphrodites/asexuals than for males and females, generalizing the results may lead to underestimating the role of outcrossing in adaptation. In other conditions, the opposite may be true, leading to the overestimation of the role of outcrossing. Thus, when investigating the role of outcrossing in adaptation to novel conditions, it is important to understand how the conditions applied influence fitness of individuals expressing particular reproductive strategies.

Many of the studies investigating the problem of male maintenance associated with the existence of outcrossing have used *Caenorhabditis elegans*, a common model species in evolutionary and genetic research (Gray and Cutter 2014). *C. elegans* is androdioecious, with hermaphrodites coexisting with males. Hermaphrodites are capable of both selfing and outcrossing with males, but they are not able to mate with other hermaphrodites (outcrossing occurs only through mating with males). Sex is determined by the ratio of X chromosomes to autosomes (Hodgkin 1987) with hermaphrodites having AA:XX genotype and males AA:X0. Hence, a male forms when a gamete carrying one X fuses with a gamete carrying no sex chromosome – which can happen either through self-fertilization following X-chromosomes nondisjunction during meiosis, or via outcrossing (since half of the gametes produced by males lack the X chromosome).

Selfing is a predominant reproductive mode in *C. elegans.* Male frequencies vary between strains (reviewed in Anderson et al. 2010). However, in most populations (including the well-studied laboratory strain N2) they are very low, often similar to those of nondisjunction events (Hodgkin 1983; Chasnov and Chow 2002; Teotònio et al. 2006). Adult males and hermaphrodites show no difference in viability (Hodkin 1987; Gems and Rddle 1996; 2000). Although males survive dauer (an alternative developmental stage induced by stressful conditions) slightly better than hermaphrodites (Morran et al. 2009), this difference is very small. Male fertilization success depends on their frequency in a population, being the highest at about 0.2: in the N2 laboratory strain males may sire 70% of the offspring produced in such populations (Stewart and Phillips 2002). However, these rates are still too low to prevent a gradual loss of males from populations. Moreover, as inbreeding depression has not been recorded in the species (Johnson and Wood 1982; Johnson and Hutchinson 1993; Chasnov and Chow 2002; Dolgin et al. 2007), suggesting that prolonged inbreeding has purged mutation load, offspring resulting from outcrossing is not predicted to be fitter than offspring of selfing hermaphrodites (cf. Anderson et al. 2010). Altogether, this suggests that males should be easily lost from populations - which is supported by the results of experiments performed under standard laboratory conditions (Steward and Plillips 2002; Chasnov and Chow 2002; Cutter et al. 2003; Cutter 2005) - and that they do not play important role in *C. elegans* evolution.

However, the fact that a large fraction of the genome is devoted to male functions (Jiang et al. 2001) and that genes expressed only in males are among the most conserved in this species (Cutter 2005) questions such reasoning. Unless *C. elegans* has become predominantly selfing only recently, male-specific genes must have been maintained and conserved by selection acting on males (Loewe and Cutter 2008). This suggests that outcrossing or/and males as such have fitness advantage in at least some conditions and circumstances. The hypothesis that outcrossing becomes favorable in populations adapting to environmental challenge has been gaining support over the last several years (Wegewitz et al. 2008; Morran et al. 2009a, b; 2013, Teotònio et al. 2012; Lopes et al. 2008; Carvalho et al. 2014, but see Theologidis et al. 2014).

Extensive knowledge about the genetics of *C. elegans* allows manipulating its mating system, providing a useful tool for experimental tests of the role of outcrossing in adaptation. Scientists have identified several mutations altering dynamics of mating systems in this species (see Anderson et al. 2010 for review), with mutations in *fog-2* and *xol-1* genes being among the most frequently used in evolutionary studies (Stewart and Phillips 2002; Katju et al. 2008; Morran et al. 2009). The first of those, *fog-2*, produces a protein inhibiting production of sperm in hermaphrodites homozygous for this locus (Schedl and Kimble 1988; Clifford et al. 2000; Nayak et al. 2005). Thus, this mutation effectively turns hermaphrodites into females, enforcing obligate outcrossing in a mutant population. Mutation in *xol-1* gene causes obligate selfing, as it disturbs dosage compensation rendering males inviable (Miller et al. 1988; Rhind et al. 1995). The possibility to utilize the above mutations enables establishment of populations differing in mating systems. This makes *C. elegans* a species in which the hypotheses considering male maintenance and the role of outcrossing can be precisely tested.

However, as mentioned above, the same change in external environment may affect fitness of males, females, and hermaphrodites differently, hence imposing disparate selective pressures on different mutants and reproductive systems. Here, we use replicated lines derived from the N2 strain to investigate the effects of a stressful novel environment (increased ambient temperature) on fitness of *fog-2* (obligatorily outcrossing) and *xol-1* (obligatorily selfing) mutants, as well as wild type, of *C. elegans.*

## Materials and methods

### Animal culture

We followed standard procedures for culturing and manipulation of *C. elegans* (Brenner 1974). Animals were grown on 6 cm Petri dishes with standard Nematode Growth Medium (NGM) seeded with 200µl of OP50 strain of *Escherichia coli* (Stiernagle 2006).

We used a wild type N2 (Bristol) strain of *C. elegans*, obtained from the *Caenorhabditis* Genetics Center (CGC, University of Minnesota, USA). In this strain, the frequency of males is approximately 0.002 (Hodkin et al. 1983; Chasnov and Chow 2002; Teotònio et al. 2006), which does not exceed the rate at which they are produced by spontaneous non-disjunction events (Hodgkin et al. 1979; Rose and Baillie 1979; Cutter and Payseur 2003; Teotònio et al. 2006). Two genetically homogeneous isolines (henceforth referred to as source lines A and B) were established by 20 generations of single hermaphrodite transfer.

### Mating system manipulations

Two reproductive system-altering mutations were independently introgressed into each of the two source lines in order to obtain two sets of obligatorily selfing, obligatorily outcrossing, and wild type (facultatively outcrossing) populations with otherwise similar genetic backgrounds.

To obtain obligatorily selfing lines, *xol-1* mutation *tm3055* was introgressed into each of the source lines (carrying the wild type *xol-1* allele, henceforth: *wt*), according to a modified protocol described by Theologidis and colleagues (2014). (1) Hermaphrodites homozygous for the *tm3055* mutation were placed on Petri dishes with an excess of source line (*wt/wt*) males (P generation). (2) The F1 hermaphrodite offspring were individually isolated and allowed to reproduce by self-fertilization for 1 day, after which they were genotyped to confirm *tm3055/wt* heterozygosity (this step was necessary since *C. elegans* hermaphrodites can reproduce by self-fertilization regardless of the presence of males, which in this case would have resulted in *tm3055/tm3055* offspring). (3) Hermaphrodite offspring (F2) of the verified heterozygotes were individually placed on Petri dishes and an excess of source line (*wt*/*wt*) males was added to each dish. (4) Their offspring (F3) were screened for the presence of males once they reached the L4 stage; the dishes containing males were discarded. The absence of males indicated that the maternal (F2) hermaphrodite was homozygous for the *tm3055* allele, making all male offspring inviable (5) From the dishes containing no males F2, hermaphrodites were individually isolated to Petri dishes and allowed to reproduce by self-fertilization for 1 day, after which they were genotyped to confirm *tm3055/wt* heterozygosity (see step (2)). Steps 2 – 5 were repeated 8 times in total.

To obtain obligatorily outcrossing lines, *fog-2* mutation *q71* was introduced into each of the source lines using a protocol described by Teotònio et al. (2012). Parental hermaphrodites from a given source line were mated with males homozygous for the *fog-2* mutation, and their hermaphrodite offspring (F1) were separately selfed to generate F2. Twenty F2 hermaphrodites from each of the lines were picked onto individual plates and selfed. F3 progeny was checked for phenotype “piano” (accumulation of unfertilized oocytes in the gonads) and absence of F4 progeny, which indicated homozygosity for *fog-2* mutation *q71* in parental hermaphrodite. There were 8 other such cycles of introgression, starting with the F3 fog-females being mated with an excess of males from the source lines.

Wild-type source lines were subjected to single hermaphrodite transfer for the time necessary to complete eight cycles of introgression in the mutant lines, before being used in the fitness assays, so the inbreeding in all lines was similar. After completing the introgression (or single transfers in wild type lines) all the lines were frozen at −80°C.

### Fitness assay

All the lines were thawed and placed on Petri dishes at 20°C. After 4 days (1 generation) of acclimatization at 20°C, the lines were synchronized and cleared of any contaminations using bleaching (Stiernagle, 2006; briefly: the procedure involves treating the animals with hypochlorite solution which dissolves adults and larvae but leaves eggs protected by shells intact). After bleaching, for each source line × breeding system combination, approximately 2000 eggs were transferred on two 14 cm diameter Petri dishes (1000 eggs per dish), one of which was subsequently placed at 20°C, and the other at 25°C. The temperature error range of the incubators was 0.5°C. The worms were left at the experimental temperatures for two generations before the fitness assay was performed. High population densities were prevented by chunking procedure, *i.e.* carving out a small piece (approx. 1 cm^2^) of agar containing worms from an old plate onto a new plate seeded with bacteria medium. The procedure was performed on the third day of each generation.

After the 2 generations of acclimation, for each source line × breeding system × temperature combination, 15 hermaphrodites (for wild type and *xol-1*) or 15 pairs (for *fog-2*) in the last larval stage (L4) were individually transferred to 6 cm Petri dishes. Each dish was then returned to its respective temperature. After 24 hours, we transferred the animals onto new plates, while the dishes with eggs were left for two days, until the offspring reached L3/L4 larval stage. We repeated this procedure (transferring adult individuals onto new plates while leaving the eggs they had laid since the previous transfer for further development) for 7 days. At the L3/L4 stage, the offspring were counted, and each scored worm was aspired out with a vacuum pump to prevent counting the same individual multiple times. Each dish was re-inspected the next day in order to score the offspring which were overlooked during the first counting (e.g. because they crawled under the agar or on the side of the dish). The total number of offspring (*i.e.,* the lifetime reproductive success) of each experimental hermaphrodite / pair was then summed up over all days it had reproduced. We eliminated from the analysis data obtained from individuals which could not be found on the plates and therefore we could not be certain when they finished reproducing.

### Statistical analyses

Data were analyzed using R.3.2.0 (R Core Team 2015).

Proportions of infertile individuals / pairs were analyzed using Fisher exact test for each of the source lines separately at each temperature. Hence, four analysis were performed, comparing infertility rates across breeding systems (analysis for the source line A at 20°C, analysis for the source line B at 20°C, analysis for the source line A at 25°C, analysis for the source line A at 25°C). Additionally we compared infertility rates across temperatures applying Fisher exact tests within each mating system and source line combination.

Data on lifetime reproductive success were first analyzed using the analysis of variance with temperature, breeding system and source line as fixed factors (including all interactions) and log-transformed number of offspring as a response variable. Log-transformation was done to reduce the heterogeneity of variances. As we found strong significant interaction between temperature and breeding system (see Results), we further applied separate analyses within each temperature and within each breeding system. These more detailed analyses were performed without transformation (*i.e.* on raw data); instead, we used gls and lme functions implemented in the nlme package to build models allowing for different variance structures in the data (Davidian and Giltinan 1995).

For each temperature, we applied analyses with breeding system, source line (both treated as fixed factors) and their interaction. Three models differing in variance structure were applied: 1) a standard ANOVA with homogenous variance structure, 2) a model allowing for differences in variances between mating systems, and 3) a model allowing for different variances between the source lines. In models 2 and 3 we used VarIdent variance structure allowing for differences in variance covariates between levels of a nominal variable (Zuur et al. 2009). Then, models 2 and 3 were compared to model 1 using log-likelihood ratio test. If both models accounting for heterogeneity of variances were significantly better than the model assuming variance homogeneity, the comparison between these two models was performed using AIC criterion (we could not perform log-liklihood ratio test as these models are not nested). The results of the model that best described variance structure are reported in the Results section.

Similarly, within each breeding system, we applied analysis with temperature, source line and their interaction. Again, we applied three models: 1) an ANOVA with homogeneous variance structure, 2) a model allowing for differences in variances between temperatures, and 3) a model allowing for different variances between the source lines. Again, in models 2 and 3 we used VarIdent variance structure (Zuur et al. 2009). Model comparisons were performed as described above.

## Results

### Infertility rates

At 20ºC, infertility rates were low and did not differ between the breeding systems (Fisher’s exact tests source line A: p=0.096, source line B: p=0.343). Two out of 14 assayed pairs were infertile in the fog line A and three out of fifteen assayed pairs were infertile in the fog line B. There were no infertile individuals in the xol lines. In the wild type lines, 0 and 2 individuals failed to lay eggs out of 15 assayed (lines A and B, respectively). In contrast, at increased temperature, the rates of infertility were strikingly high in the fog lines; 10 and 7 out of 15 tested pairs were infertile in the fog lines compared to 2 and 0 out of 15 in the xol lines and 1 and 0 out of 15 in wild type lines (lines A and B, respectively; Fisher’s exact test, p<0.001 for both line A and line B).

Fertility rates decreased significantly with temperature in the fog line A (Fisher’s exact test, p=0.008). The fog line B also tended to show reduction in fertility rate, although the trend was not significant (p=0.109). Fertility did not vary between temperatures in the xol lines (Fisher’s exact test, line A: p=0.483, line B: p=1) or the wild type lines (Fisher’s exact test, line A: p=1, line B: p=0.483).

### Lifetime reproductive success

Analysis of variance performed on log-transformed data showed significant influence of temperature (F1,167=32.374, p<0.001) and breeding system on lifetime reproductive success (F2,167= 26.396, p<0.001), but no significant effect of source line (F1,167= 0.029, p=0.865). There was a significant interaction between breeding system and temperature (F2,167=16.149, p<0.001; Fig. 1). Therefore, we performed separate analyses within temperatures and mating systems.

**Figure 1.**
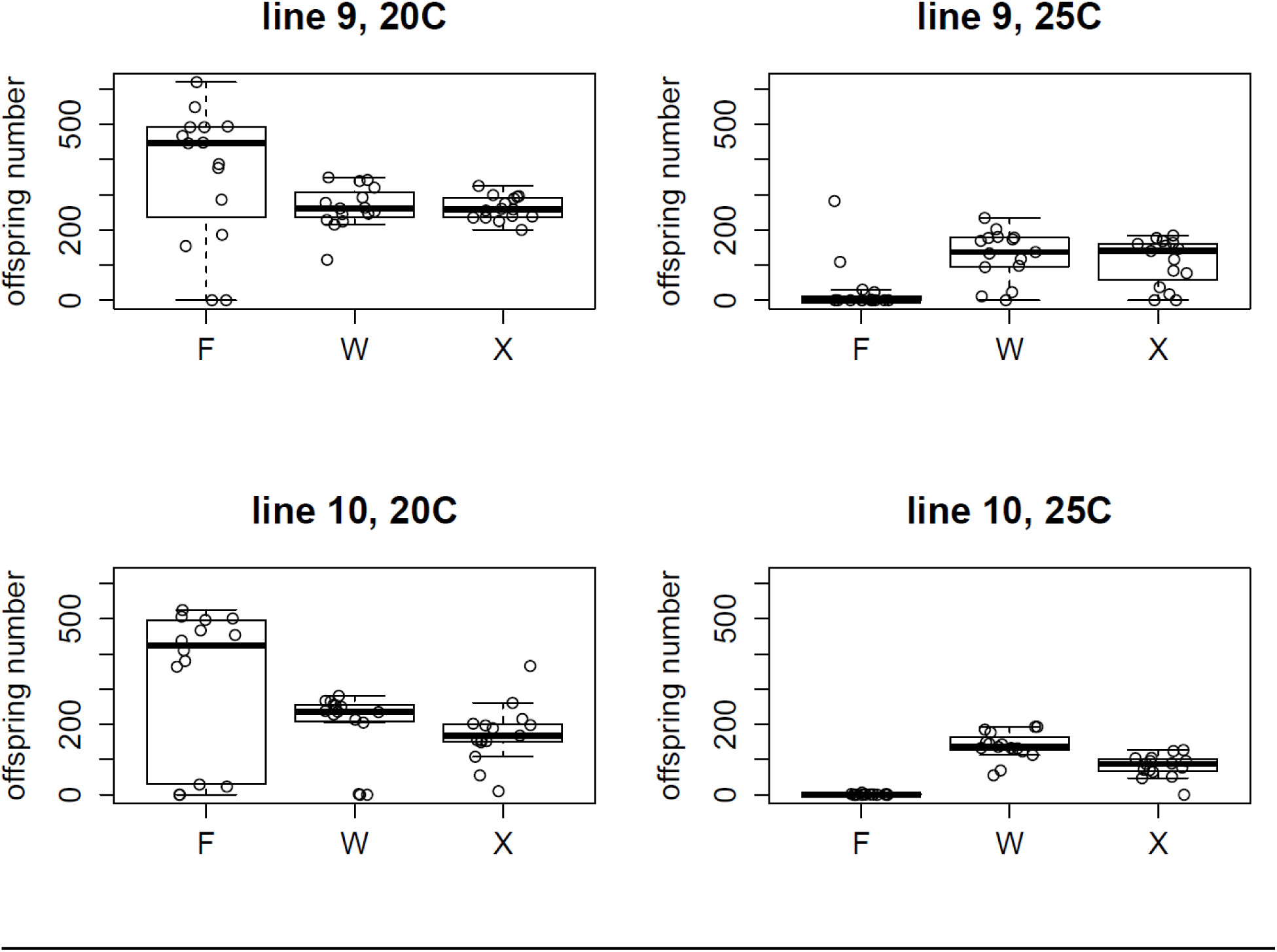
Boxplot diagrams of the number of offspring produced by the fog (F), wild type (W) and xol (X) lines for each of the source lines and thermal treatments. The boxes signify the upper and lower quartiles, and the median is represented by a solid line.

At 20ºC, the model allowing for the differences in variances between the breeding systems fitted our data significantly better than the model with homogeneous variances (Log-likelihood ratio L=41.594, df=2, p<0.001). The model with different variances between the lines was not better than the homogeneous variances model (L=1.376, df=1, p=0.241). Therefore, we report the results of the model allowing for variance differences between breeding systems.

Lifetime reproductive success differed significantly between breeding systems (F2,83= 5.285, p=0.007) with fog lines being the most fecund (Fig. 1) and between source lines (F1,83= 19.620, p<0.001); the interaction between breeding system and source line was not significant (F2,83=0.354, p=0.703).

At 25ºC, the model with variances differing between source lines described variance structure of our data most properly (comparison with the homogeneous variance model: L=28.586, df=1, p<0.001). The model with variances differing between breeding systems was not better than than the model with homogeneous variances (L=0.309, df=2, p=0.857).

Lifetime reproductive success at 25ºC differed between breeding systems (F2,84 =87.316, p<0.001), with fog lines producing the lowest number of offspring (Fig.1), but not between source lines (F1,84 =1.779, p=0.183). Also, breeding system by source line interaction was insignificant (F2,84=1.232, p=0.297).

Analyses within mating systems showed decrease of lifetime reproductive success at increased temperature in the fog lines (model with variances differing between temperatures, F1,55 =71.897, p<0.001) with no significant difference between source lines (F1,55 = 2.354, p=0.131) or source line by temperature interaction (F1,55 =0.001, p=0.970). Lifetime reproductive success of the xol lines differed between temperatures (homogeneous variances model, F1,56= 66.444, p<0.001) and source lines (F1,56=15.246, p<0.001). There was also a significant influence of the interaction between temperature and the source line (F1,56=4.267, p=0.044). In the control (wild type) lines, lifetime reproductive success also decreased at increased temperature (model with different variances between the source lines, F1,56=26.664, p<0.001). There was a significant interaction effect between temperature treatment and source line (F1,56= 4.519, p=0.038) with insignificant source line effect (F1,56= 0.767, p=0.385).

## Discussion

Our study reveals the reduction of reproductive success at high temperature in all three breeding systems. This is in line with previous studies demonstrating that *C. elegans* fecundity is highest at 20°C and declines with increasing temperature (e.g. Byerly et al. 1976; McMullen et al. 2012; Petrella 2014). The decrease of reproductive function at high temperatures seems to be associated with functioning of both spermathogenic and oogenic germ lines (Aprison and Ruvinsky 2014; Petrella 2014), although the relative contribution of these two factors varies between strains (Petrella 2014).

More interestingly, the influence of temperature on reproductive success was much more dramatic in the outcrossing (fog) lines compared to both xol and wild type selfing lines (Fig. 1). In particular, the proportion of pairs that did not produce any eggs rose from 0.13 and 0.14 to 0.67 and 0.47 in the fog lines. In contrast, in selfing lines, while the offspring number declined with increased temperature, the infertility rates remained low (xol: 0.13 and 0.07, wild type: 0.07 and 0). The decrease of fertility was only significant in one of the fog lines. However, in the other outcrossing line a clear trend in the same direction was observed together with extremely low numbers of offspring produced by fertile pairs (only one or two offspring produced by all the fertile pairs).

Such drastic fitness decline in outcrossers under thermal stress could have resulted from male and/or female gamete production failure. Indeed, it has been shown that increased temperature affects sperm and oocyte production, ovulation and spermatid activation in *C. elegans* (Aprison and Ruvinsky 2014; Petrella 2014). We might also expect thermal stress to result in elevated gamete death (McMullen et al 2012) as increased temperature during ovulation has been shown to reduce gamete viability in some fish species (Van Der Kraak and Pankhurst 1997).

However, low incidence of sterility in selfing lines prove that both types of gametes successfully function in hermaphrodites under the same thermal stress. Hence, we hypothesize that the higher thermal sensitivity of obligatorily outcrossing lines may be associated with mating failure.

Temperature is well-known to affect behavior of ectotherms (reviewed by Angiletta 2009) including reproductive behavior in many species (Wilkes 1963; Linn and Campbell 1988; Katsuki and Miyatake 2009). Whereas self-fertilization is a purely physiological process, outcrossing requires a complex set of behaviors in *C. elegans*. First, males have to respond to chemosensory cues from potential partners (Simon and Sternberg 2002). Then, they need to locate the vulva, to which they insert their spicules and ejaculate (Barr and Garcia 2006). Such a complex process is likely to be sensitive to environmental conditions, as disturbance at any of its components will result in reduced mating ability of an animal. We are not aware of any studies specifically addressing thermal effects on *C. elegans* mating behavior, however, it has been shown to be sensitive to intrinsic stress caused by senescence. Chatterjee et al. (2013) demonstrated that reproductive senescence in *C. elegans* males is associated with decreased mating efficiency rather than deterioration of sperm quality or sperm number.

Elucidating the mechanisms behind the pattern observed in our study requires further work. Whatever the mechanism, however, our results highlight the fact that the level of stress created by the same change in external environment (5 ºC increase in ambient temperature) can differ dramatically between reproductive systems. Importantly, the sharp decline in mean fitness of outcrossing lines was also associated with an interesting pattern of variation: while 57% of pairs were unfertile and 30% only produced 1-6 larvae, four pairs (13%) bred 23, 30, 109 and 282 offspring, respectively. In hermaphrodites, such a heterogeneous response has been observed only under much more severe thermal stress (≥ 28 °C; Mc Mullen et al. 2012).

Such a pattern of response to a stressful environmental factor can have complex effects on adaptation process in outcrossing populations. On one hand, very low (or zero) fitness of most individuals translates to low effective population size, which increases the impact of genetic drift and may also lead to inbreeding depression, hampering adaptive potential and increasing the risk of extinction. On the other hand, large variation in fitness will generate strong selection on any traits associated with it (Lynch and Walsh 1998) which can increase the rate of adaptation. Furthermore, if mating efficiency does indeed strongly contribute to the reproductive performance of outcrossers, as we hypothesize, high temperature will impose strong selection on traits associated with mating success. This would further lead to intense sexual selection over mating, making sexual selection an important contributor to adaptation process in populations of outcrossers (Candolin and Hauschele 2008; Lorch et al. 2003; Plesnar-Bielak et al. 2012).

Our results have important implications for investigating the evolution of reproductive modes in the context of environmental changes. They suggest that the relationship between reproductive system and the dynamics of adaptation to environmental changes (Anderson et al. 2010) may be driven not only by the level of genetic shuffling, but also by differences in selective pressures these changes impose on different reproductive modes. As these differences are likely to depend on the nature of the environmental change, the conclusions drawn from experiments subjecting outcrossing and selfing populations to novel conditions may therefore be limited to the specific environmental manipulation applied. Thus, to correctly interpret the results of such studies and, more importantly, to be able to estimate to what extent they could be generalized, it is important to know how different environments influence fitness of outcrossers and selfers. Therefore, a logical next step would be to test how hermaphrodite, male and female fitness is influenced by a variety of different stressors.

In conclusion, our results show that the same environment may impose different selective pressure depending on the reproductive system in a given population. This can have substantial effects on the way in which a population evolves in response to this factor. All this should be taken into account when empirically assessing the role of males and outcrossing in adaptation process.

## Acknowledgements

The study was supported by National Science Centre, grant UMO-2013/09/B/NZ8/03317for Z.M.P.

